# Organ-on-a-chip model of vascularized human bone marrow niches

**DOI:** 10.1101/2020.04.17.039339

**Authors:** Drew E. Glaser, Matthew B. Curtis, Peter A. Sariano, Zachary A. Rollins, Bhupinder S. Shergill, Aravind Anand, Alyssa M. Deely, Venktesh S. Shirure, Leif Anderson, Jeremy M. Lowen, Natalie R. Ng, Katherine Weilbaecher, Daniel C. Link, Steven C. George

**Affiliations:** Department of Biomedical Engineering, University of California, Davis; Department of Chemical Engineering, University of California, Davis, Davis, CA 95616; Department of Biomedical Engineering, Washington Univeristy in St. Louis; Department of Medicine, Washington University in St. Louis, St. Louis, MO 63130

## Abstract

Animal models of bone marrow have limited spatial and temporal resolution to observe biological events (intravasation and cellular egress) and are inadequate to dissect dynamic events at the niche level (100 microns). Utilizing microfluidic and stem cell technology, we present a 3D *in vitro* model of human bone marrow that contains perivascular and endosteal niches complete with dynamic, perfusable vascular networks. We demonstrate that our model can perform *in vivo* functions including maintenance and differentiation of CD34^+^ hematopoietic stem/progenitor cells (HSPC) for up to fourteen days, egress of myeloid progenitors, and expression of markers consistent with *in vivo* human bone marrow. The platform design enables the addition of tissue niches at a later timepoint to probe mechanisms such as tumor cell migration. Overall, we present a novel organ-on-a-chip platform that is capable of recapitulating the human bone marrow microenvironment to observe hematopoietic phenomena at high spatial and temporal resolution.

## Introduction

The bone marrow is comprised of many cell types and has a unique sinusoidal blood supply that gives rise to distinct microenvironments or niches (*1, 2*) that are small in size (order 100 microns) and are in close proximity. The perivascular and endosteal niches are of particular interest due to their support of hematopoiesis (*3, 4*), a primary function of bone marrow, but also their impact on disease processes such as tumor metastasis and dormancy (*1, 5, 6*). The endosteal niche is thought to be comprised of osteoblast cell at the surface of bone where a hematopoietic stem cell (HSC) resides, but may include the adjacent 50 μm of bone (*7, 8*). Similarly, the space immediately adjacent to the abluminal surface of a blood vessel comprises the perivascular niche (*4, 9*).

Both the perivascular and endosteal niches contain sinusoidal blood vessels as a key feature of the bone marrow. These specialized vessels are surrounded by many different cell types including mesenchymal stem cells (MSC), specialized CXCL12 abundant reticular (CAR) cells, HSC, leukocytes at different stages of differentiation, and adipocytes (*10*). Bone marrow stromal cells are known to secrete chemotactic signals such as stem cell factor (SCF) (*4*) and CXCL-12/SDF-1 (*11*). Endothelial cells express specialized ligands on the luminal surface such as E-selectin, ICAM, and VCAM (*3, 12*). These chemokines and adhesion molecules cooperatively work to facilitate homing of circulating HSC to bone marrow (*3, 13*). *In vivo,* osteoblasts also express VCAM and CXCL-12 in addition to osteopontin, which has been implicated in maintaining HSC quiescence (*7, 14*). Moreover, blood vessels found within the endosteal niche express higher levels of E-selectin, which play an important role in regulating HSC cell cycling (*15*), and has been implicated as a regulator of breast cancer metastasis to bone (*12, 16*). We endeavored to create a unified model of these two bone marrow niches (perivascular and endosteal) using an *in vitro* using a microfluidic organ-on-a-chip system.

Advances in tissue engineering and microfluidics have created “organ-on-a-chip” technologies to generate 3D microphysiological mimics that utilize a broad range of human cells, thus providing specific advantages over traditional 2D culture and mouse models. Moreover, microfluidic technology lends itself well to capturing the spatial scale of the adjacent endosteal and perivascular bone marrow microenvironments. Although several labs have utilized similar tissue engineering and microfluidic approaches to recreate aspects of the perivascular and endosteal niches (*1, 5, 17, 18*), however, these models have lacked vital aspects of *in vivo* tissue. Our model incorporates independent spatial and temporal control of the niches, a vascular network that is essential for niche maintenance and function (*4, 19*), and the ability to maintain a population of multipotent HSPC. Herein, we present a microfluidic-based platform that mimics all of these important features of the bone marrow microenvironment; namely, a 3D dynamic vascular network, the perivascular and endosteal niches, and maintenance and differentiation of CD34+ HSPC.

## Results

### Device Design Recreates Scale and Physical Environment of the Bone Marrow

The device was designed to create the endosteal and perivascular microenvironments in the bone marrow while maintaining it’s fundamental biological function - hematopoiesis. To this end, the symmetrical two-way port between the chambers was designed to enable communication and cell trafficking between the adjacent chambers (Fig. 1C).

**Figure 1.**
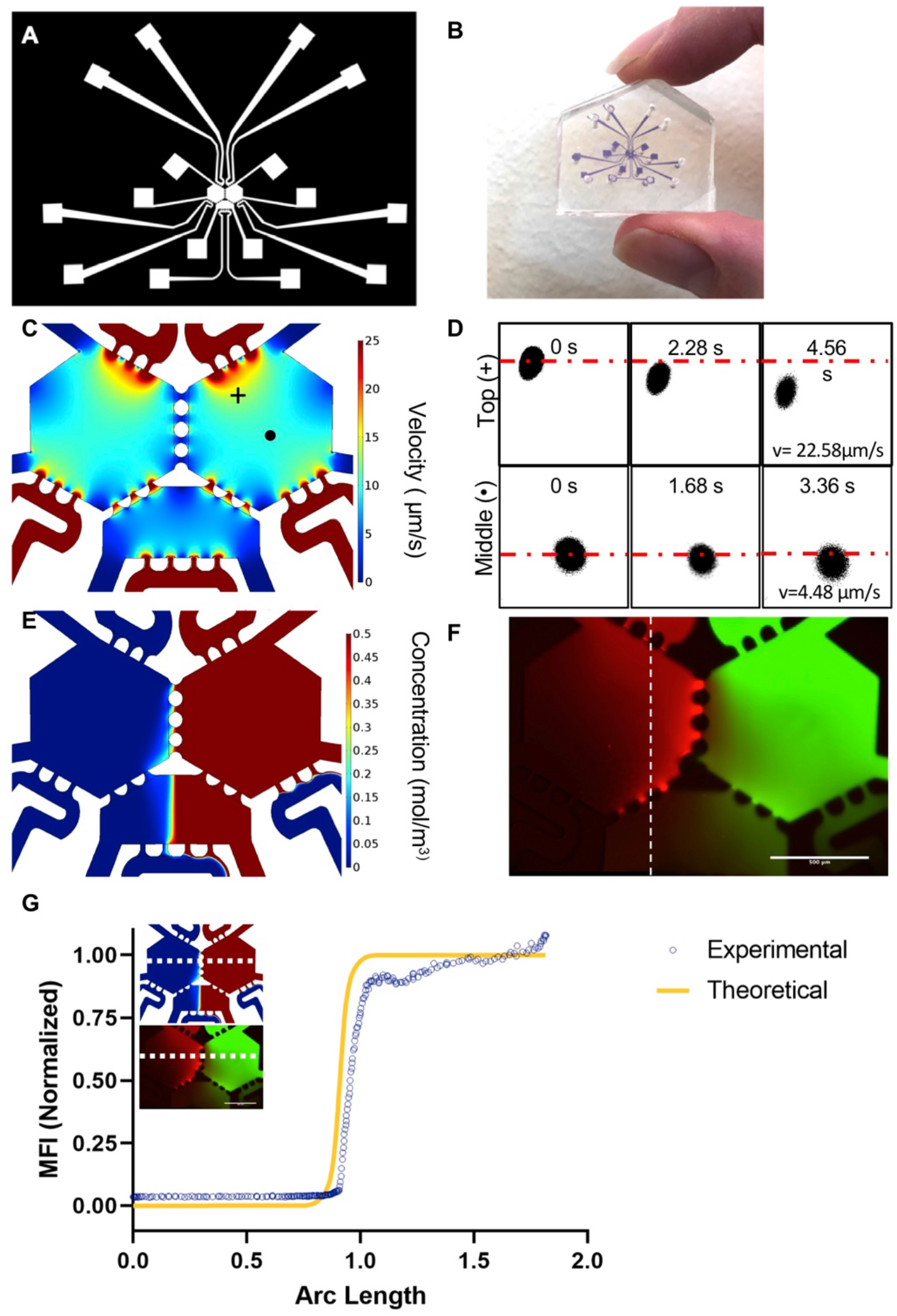
Characterization of transport phenomena in a three chambered-microfluidic device. (**A**) CAD design depicts the overall device design with connecting fluidic lines. (**B**) Actual size of the device. (**C**) COMSOL simulation of interstitial flow through the fluidic device predicts flow rates of ~10 μm/s. (**D**) FRAP experiments in the top and middle of the device show an interstitial flow velocity of 22.58 μm/s and 4.48 μm/s respectively. Average interstitial flow was 17.7 μm/s, n=3. (**E**) COMSOL modelling of 70 kDa dextran through the right side of the device predicts limited diffusion of dextran through the ports. (**F**) 70 kDA TRITC and FITC dextran perfused through a 10 mg/mL fibrin gel in the device shows limited diffusion from one chamber to another and an even distribution of dextran in the bottom central chamber. (**G**) Mean fluorescent intensity of an arbitrary line (white dots) through the middle ports of the simulated device and the actual device correlate with each other across the distance of the device.

Our lab has previously demonstrated that interstitial flow plays a critical role in the formation of dynamic, perfusable microvascular networks in microfluidic devices by vasculogenesis (*23, 32*). Therefore, as a first step, the interstitial flow was modelled in COMSOL to identify conditions that were favorable for vascular formation (*20, 23*). Pressure heads in a single device of 20, 18, 10, 11, 5, and 4 mm H_2_O (Fig. S1) were used to simulate interstitial flow in the device. Using these pressures, interstitial flow rates ranged from 6-20 μm/s depending on position in the chamber(s), with slower flow rates in the middle of the chamber and higher flow rates near the ports (Fig. 1C and S1). This flow rate has been shown to support capillary formation in microfluidic devices by vasculogenesis (*23, 32*). Theoretical interstitial flows were verified experimentally using FRAP with an Olympus FV3000 confocal microscope (Fig. 1D). Using the previously described pressure heads, an experimental interstitial velocity of 17.7 μm/s (± 4.71 μm/s, n =3) was measured adjacent to the high-pressure ports as well as a lower velocity of 4.48 μm/s (n=1) in the middle of the device. These values align with the theoretical values measured in the same regions.

The transport of molecules similar in size to secreted factors was also modelled in COMSOL (Fig. 1E) and validated experimentally using 70 kDA dextran labelled with TRITC and FITC (Fig. 1F). Because of the symmetric design of the chambers and the operating conditions, convective flow between the chambers is near zero (Fig. 1E, F), and communication is limited to diffusion. This convective isolation also occurs in the adjacent bottom chamber, with a sharp spatial transition (concentration gradient) in the middle of the chamber (Fig. 1E-G). The mean fluorescent intensity of the FITC dextran in the bottom chamber (dashed line in Fig. 1E, F) mapped closely to the theoretical concentration profile (Fig. 1G).

### BMoaC Recapitulates *in vivo* markers of BM

To generate the BMoaC, EC were mixed with hOB or BMSC in a fibrin hydrogel and injected into either the left or right chamber, respectively, to generate a tissue (Fig. 2A). Over a period of 4 to 7 days, vascular networks were apparent within both niches (Fig. 2B-H, S2A). At 7 days, some devices were fixed and stained for transmembrane proteins Stem Cell Factor (SCF), ICAM-1 (CD54), VCAM-1 (CD106), and E-selectin (CD62E) as well as the basement membrane marker, laminin (Fig. 2B-F). EC were counterstained for either CD34 or CD31. EC expressed SCF-1 on the abluminal side of the vessel wall in both niches (Fig. 2Bi,ii). hOB also stained for SCF throughout the niche as well some BMSC. The basement membrane protein laminin was clearly seen on the abluminal side of the vessel wall in both niches (Fig. 2Ci,ii). The cellular adhesion markers ICAM-1, VCAM-1, and E-Selectin were also expressed by EC in both niches (Fig 2. D-F) without stimulation of inflammatory factors such as TNFα. Co-expression of these markers with an EC-specific marker (CD34 or CD31) was easily detected at high magnification (Fig. 2Di-Fii). Osteoblasts also expressed VCAM-1. Vascular networks were positive for nestin (Fig. 2G), which has been shown to identify arterioles in the bone marrow (*3*). Vascular networks, formed with these stromal cells, were perfusable (Fig 2H., Sup. Vid. 1), and had similar indices of vessel network morphology (e.g., area and numbers of junctions (Fig. S2B)). The permeability of the endosteal and perivascular networks to 70 kDa dextran was 17.0 ×10^−7^ cm/s (± 8.90 ×10^−7^cm/s) and 5.2 ×10^−7^ cm/s (± 3.82 ×10^−7^ cm/s), respectively, which were statistically different (p <0.05). Vascular networks formed with osteoblasts had significantly shorter total and average vessel length (Student’s t-test p < 0.05; S2B) and a higher number of vessel endpoints (p<0.001) compared to the matched perivascular networks formed with BMSC.

**Figure 2.**
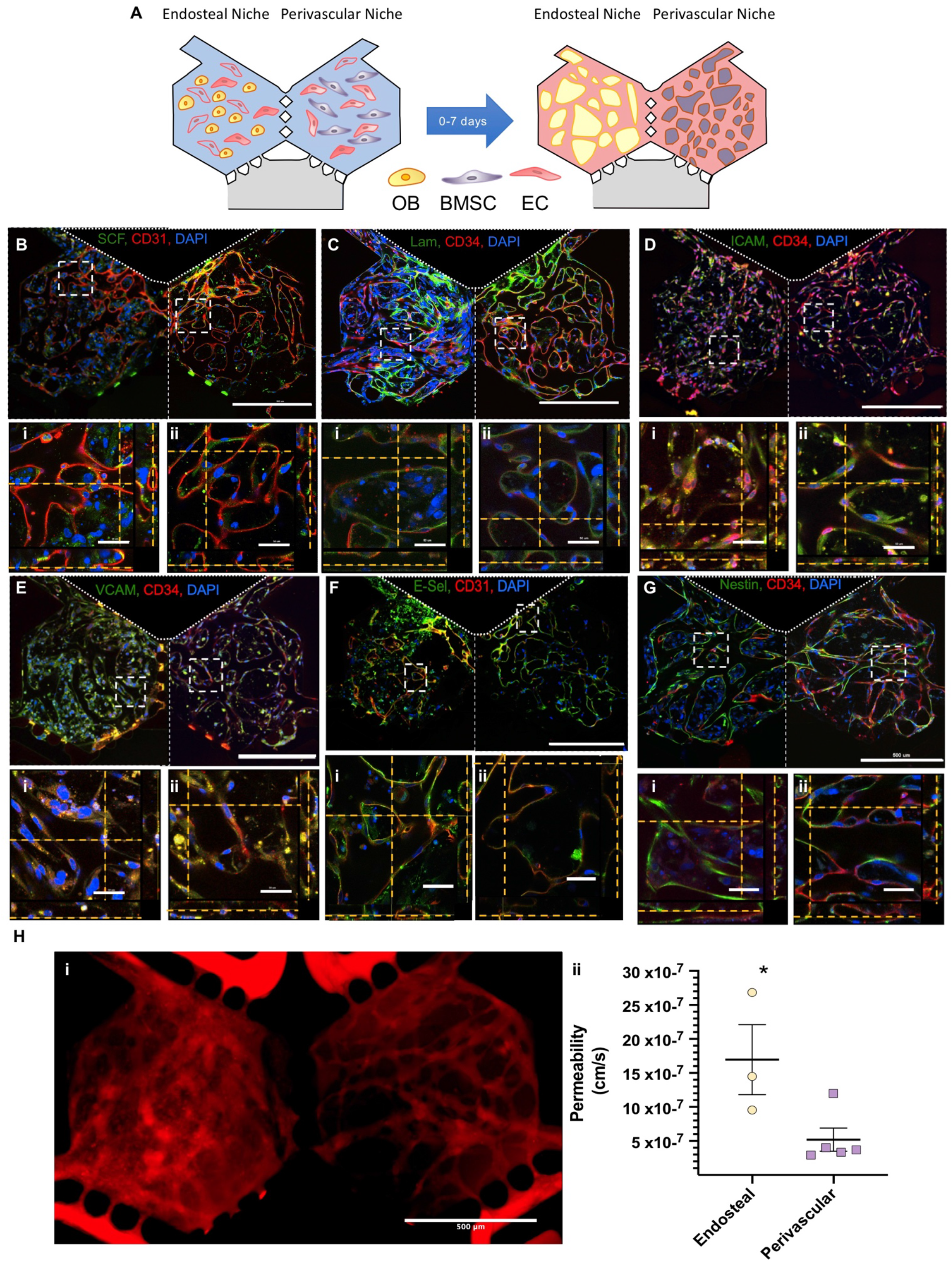
ECFC-EC in BMoaC express proteins found in native bone marrow. (**A**) Cartoon schematic of experimental design. ECFC-EC seeded with either hOB (Endosteal Niche) or BMSC (Perivascular niche) form microvascular networks in a period of 4 to 7 days. (**B**)-(**G**) EC (red, CD31 or CD34) express (**B**) SCF, (**C**) Laminin, (**D**) I-CAM, (**E**) V-CAM, (**F**) E-Selectin/CD62E and (**G**) Nestin. ROIs show z-stacks in either the (**i**) endosteal or (**ii**) perivascular niche. Nuclei counterstained with DAPI (Blue); Scale bars are 500 μm or 50 μm for high magnification view. (**H**) BMoaC microvascular networks are i) perfused with 70 kDA TRITC dextran and ii) have permeabilities of 16.96 × 10^−7^ cm/s and 5.19 × 10^−7^ cm/s for the endosteal (n=3) and perivascular niches (n=5), respectively.

Both niches stained positive for osteopontin (Fig. 3Ai-ii). Osteopontin^+^ cells in the endosteal niche were not observed adjacent to vessel walls but were observed in close association with EC in the perivascular niche. Both niches also stained positively for SDF-1/CXCL-12 (Fig. 3C, i-ii), although the expression was visibly higher in the stromal cells and some EC within the perivascular niche (Fig. 3B, ii). Very few stromal cells were leptin^+^ in the endosteal and perivascular niches (Fig. 3C, i-ii). Stromal cells were characterized as individual populations by flow cytometry. Both populations of stromal cells had extremely low levels of CD31 (<0.1%), CD34 (<5%), and CD45 (−0.05%), CD36 (~1.4%), and CXCL-12 (~3%) while expressing similar levels of CD44 (~96%) and CD90 (~96%) (Fig. 3D), as assessed by flow cytometry. The hOB and BMSC also expressed the following respective markers, albeit at different levels: CD105 (98 vs 65%), CD73 (98 vs 76%), NG2 (~49 vs 97%), CD10 (83 vs 52%), and ALP (22% vs 58).

**Figure 3.**
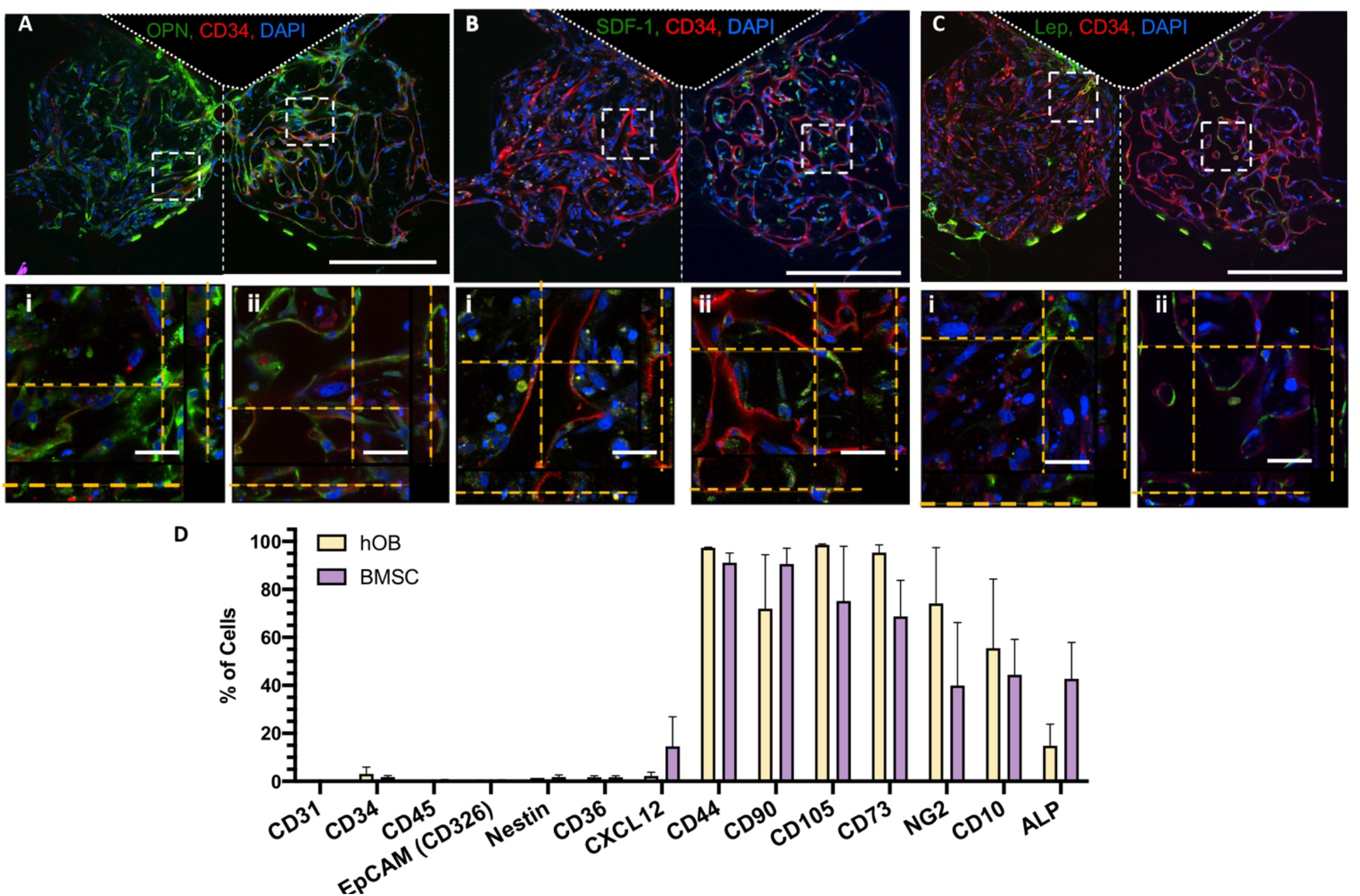
Stromal cells cultured in BMoaC express proteins found in native marrow. (**A**) Osteopontin (green) is expressed in the (**i**) endosteal and (**ii**) perivascular niches. EC (red) counterstained with CD34. (**B**) SDF-1 is detected adjacent to EC (red) in the (**i**) endosteal niche with more stromal cells appearing to express SDF-1 in the (**ii**) perivascular niche. (**C**) EC (red) are adjacent to stromal cells that express leptin as seen in the (i) endosteal and (ii) perivascular niche. Nuclei counterstained with DAPI (blue) (D) Marker expression of hOB and BMSC as determined by flow cytometry, n=3.

### BMoaC Supports Maintenance of Hematopoietic Stem/Progenitor Cells

To test whether BMoaC could support the maintenance of a small population of hematopoietic stem/progenitor (HSPC) cells, CD34^+^ cells isolated from cord blood were cultured in the device over a period of 7 to 14 days (Fig. 4A). Individual experiments were carried out with a single donor (eight donors total) and purity of the sample was assessed by flow cytometry before loading. Cord blood cells were on average 89.8% ± 2.2 CD34^+^ and 76.4% ± 2.5 CD34^+^/CD133^+^; samples with purity below 80% CD34^+^ cells were not used (Fig. 4B, S4.). There was an average of 65.6 ± 8.4 and 35.4 ± 3.0 CD34^+^ cells loaded into the endosteal and perivascular chambers, respectively, which was statistically different by t-test (Fig. 4B(ii), p < 0.05). CD34^+^ derived cells were visible by brightfield in both niches as early as day 4 and proliferated within the device over the culture period (Fig. S3B). Small round cells could be clearly seen within the lumen of blood vessels in the perivascular niche (Fig. S3B, Day 12). CD34-derived cells were also observed trafficking within the main chambers of the BMoaC as well as into the fluidic lines using time-lapse microscopy (Sup. Vid. 2). CD34^+^ cells loaded into the device without EC or stromal cells failed to proliferate after 7 days.

**Figure 4.**
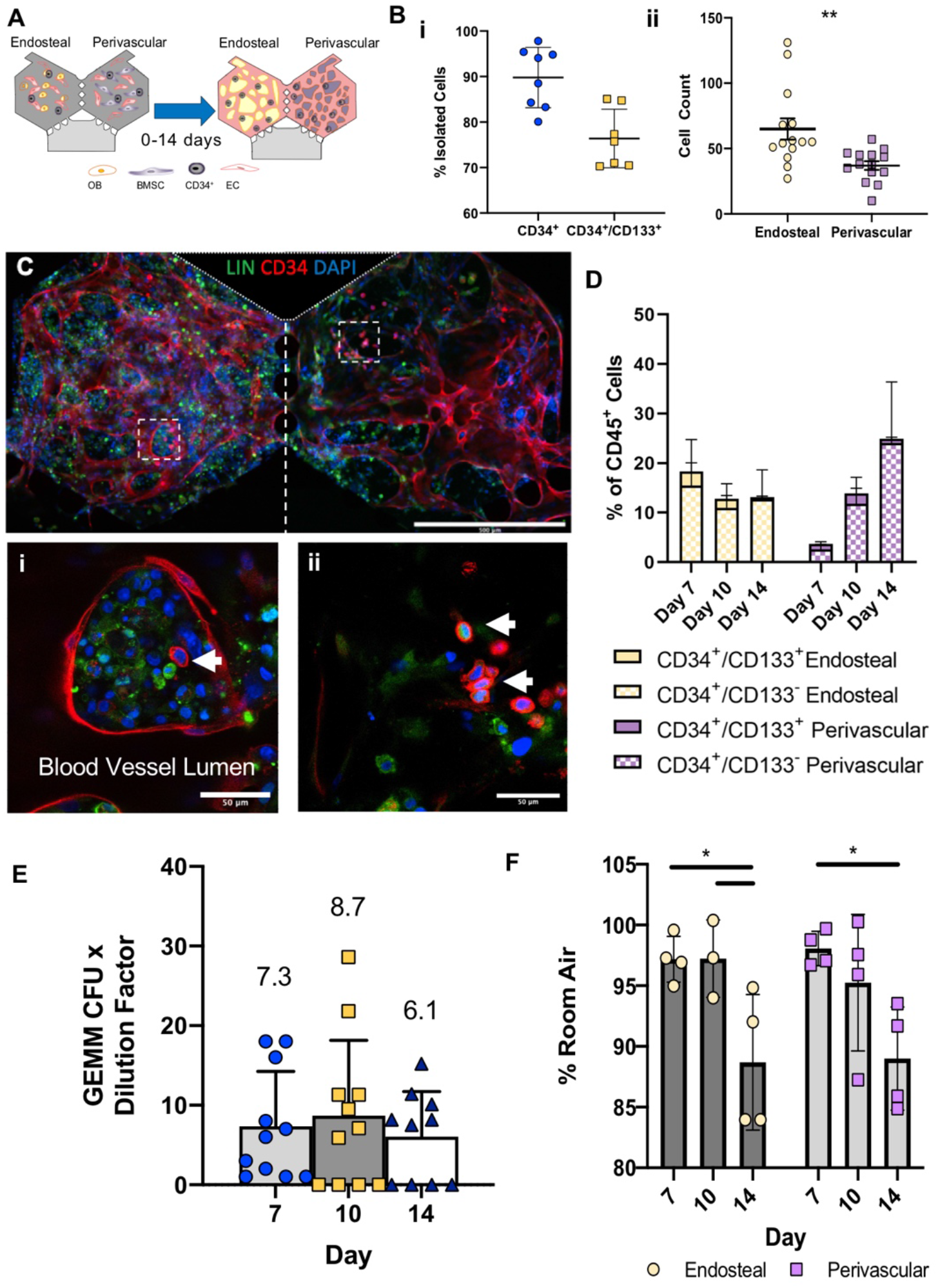
BMoaC maintains a population of CD34+ HSPC. (**A**) Experimental design. (**B**) (**i**) CD34+ cells isolated from human cord blood are 89.8% ± 6.6 CD34+ and 76.4% ± 6.4 CD34+/CD133+, b=8. (**ii**) ~100 CD34+ HSPC are loaded into each device, with ~65 and ~37 CD34+ HSPC loaded into the endosteal perivascular niches, respectively, n=13 devices, unpaired t-test **p<0.01 (**C**) BMoaC supports CD34+ blood vessels as well as (i,ii) small, round CD34+ HSPC are present along with Lin+ cells. Scale bars: 500 μm and 50 μm (**D**) Cells isolated after day 7, 10, and 14 days of culture in the BMoaC were analyzed for CD34/CD133 from both the endosteal and perivascular niches, n=5. (**E**) Cells isolated after day 7, 10, and 14 days of culture in the BMoaC consistently generate GEMM-CFU over time, n=10-11. (**F**) Bar graph of mean endosteal and perivascular O_2_ tension over time with individual data points represented by beige circles (endosteal) or purple squares (Perivascular), n=4. * represents significant difference as calculated by a 2way ANOVA, α=0.05.

To test whether a small population of CD34^+^ cells were maintained, BMoaC were either analyzed with confocal microscopy or the matrix digested and cells released to be assessed by flow cytometry for CD34, CD133, and lineage (Fig. S3C-D). Some devices contained clusters of Lin^+^ cells with kidney-shaped/multi-lobular nuclei as well as small, round CD34^+^ cells surrounded by a blood vessel in both niches (Fig. 4Ci, S3D, Sup. Vid. 3). BMoaC were able to maintain a population of Lin^−^/CD34^+^ cells as well as a differentiated population of Lin^+^/CD34^−^ cells, as identified by flow (Fig. S3D). Both niches were able to maintain a small CD34^+^/CD133^+^ population (Fig. 4D) over the culture period. Devices loaded with CD34^+^ HSPC alone in a fibrin gel were not viable. The endosteal did not see a significant change in either the CD34^+^/CD133^+^ or CD34^+^/CD133^−^ population of HSPC after day 7 (n=5). While changes in the population percentages were not significant (n=5), there was an upward trend in the amount of these stem cell-like cells that were present within the perivascular niche.

Methocult assays were performed with cells isolated from chamber digests to assess the colony forming unit (CFU) potential of the cells. Cells isolated from the devices after 7, 10, and 14 days of culture were capable of forming CFU-GEMM (Fig. S4), or multipotent colonies representing granulocyte, erythroid, monocyte, and megakaryocyte lineages (Fig. 4E). The number of HSPC on each day and in each device may be estimated from the number of CFU-GEMM colonies and the number of CD45^+^ cells isolated per device. The number of cells capable of generating a CFU-GEMM colony was stable between day 7 and day 14 (7.36 ± 6.7 and 6.0 ± 5.6 cells per device, respectively; p > 0.05 by one-way ANOVA).

Endosteal and perivascular niche devices were incubated at 37°C room air for 14 days. Mean O_2_ tension within each niche was determined using PhLIM on days 7, 10 and 14 (Fig. S5). O_2_ tension at day 14 (88.7% of room air) was slightly lower than at day 7 (97.5%) (Fig. 4F) within both niches, consistent with enhanced cellular density and metabolism and clearly demonstrating adequate oxygenation.

### BMoaC Support CD45^+^ Cell Differentiation and Expansion

BMoaC were able to support the differentiation of CD34^+^ cells into Lin^+^ cells over time. Bone marrow generates leukocytes that participate in the innate immune system; as such, we looked for the presence of CD14, a marker commonly found on monocytes. Both niches contained CD14^+^ cells in extravascular spaces (Fig. 5A). A few of these CD14+ were found within the blood vessels (Fig. 5Ai-ii), with additional CD14^+^ cells adjacent to the vessels. We next wanted to verify whether the cells that exited the tissue chamber and entered the fluidic line also showed mature markers of differentiation. To this end, we stimulated devices with M-CSF and IL-3. Many cells were observed in the fluidic lines of devices from day 6 onward and were collected through day 8 for additional culture in a 24-well plate, to achieve an adequate number of cells for flow cytometry. After 14 days of total culture, 80.3% of cells were CD15^+^CD33^+^ (consistent with a neutrophil, Fig. 5B-C).

**Figure 5.**
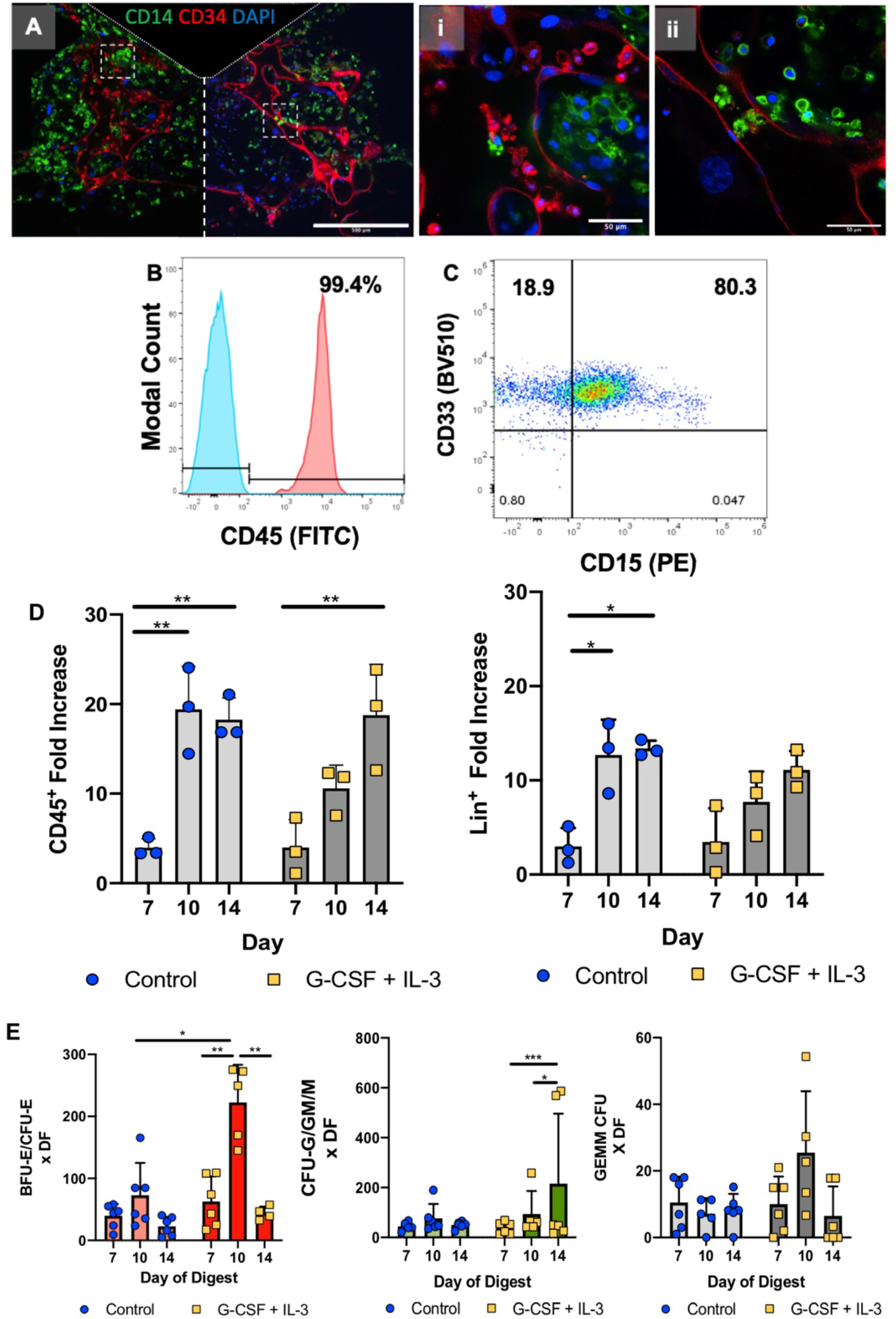
BMoaC support cellular expression of markers associated with the myeloid lineage. (**A**) After 10 days of culture, BMoaC contain cells expressing CD14 (Green) present in both sides of the device as well as in the (i, ii) lumen of CD34^+^ blood vessels (red), nuclei were counterstained with DAPI (blue). (**B, C**) Cells that left a through the fluidic line of stimulated devices were collected between day 8-14 and analyzed for the leukocyte common antigen (**B**) CD45 and the (**C**) myeloid marker CD33 and neutrophil marker CD15. Devices were stimulated with 30 ng/mL each of GM-CSF and IL-3 from Day 4-14, p<0.05 by ANOVA. (**D**) CD45^+^ cells increased from 4-20 fold; Lin^+^ cells increased 4-15 fold over the 14 day culture period compared to starting numbers of CD34^+^ HSPC, n=3 (**E**) Cells isolated after day 7, 10, and 14 days of culture in the BMoaC were capable of forming BFU-E/CFU-E, CFU-G/GM/M, and CFU-GEMM colonies, n=4-6. *p<0.05, **p<0.01, ***p<0.001 by 2way ANOVA.

Devices were also stimulated with G-CSF and IL-3 after 4 days of culture for an additional 10 days. Within each condition, there was a significant increase (p <0.01) in the CD45^+^ population of cells between day 7 and day 10, and between day 7 and day 14 for the devices stimulated with G-CSF and IL-3 (Fig. 5D). While no significant differences were found between control and stimulated devices, the total number of Lin^+^ cells within the BMoaC significantly increased from ~3-fold on day 7 up to ~12-fold on day 14 compared to Day 0 (Fig. 5D). These numbers do not include the cells that entered into the fluidic lines of the device and are analogous to a circulating progenitor cell.

We performed a methocult assay to determine the number of committed erythroid progenitors that could form BFU-E or CFU-E type colonies as well as myeloid progenitors that formed CFU-G, CFU-M, CFU-GM colonies. We did not discriminate between the different types of erythroid or myeloid colonies. Colonies formed by erythroid progenitors were the most common colonies, followed closely by colonies formed from myeloid progenitors (Fig. S4). As with the CFU-GEMM assays, we estimated the number of CFU colonies over time based on the increased population of CD45 cells. There was no significant difference found between the total number of colonies formed when devices were treated with G-CSF and IL-3 compared to the control (Fig. S4). There was a significant increase in the number of CFU-E/BFU-E colonies formed from day 10 digests when devices were treated with G-CSF and IL-3, compared to the control (Fig. 5E, p < 0.05). Cells isolated on day 10 from devices treated with G-CSF and IL-3 generated significantly more colonies than those on day 7 or 14 (Fig. 5E, p <0.01). For the devices treated with G-CSF + IL-3, there was a significant difference between day 7 and 14 as well as day 10 and 14 (p <0.05) for the number of cells capable of forming myeloid colonies (Fig. 5E). There was no significant difference between the number of GEMM colonies that formed from cells in devices digested between day 7 and 14, which was consistent with previous observations (Fig. 5E).

### Breast Cancer Cells Migrate into BMoaC Niches

A third chamber was included in the device design to observe migration of an additional cell type of interest (e.g., cancer cells) into the vascularized bone marrow chambers. To demonstrate this feature, devices were cultured for 4 days to enable development of the niche before either RFP MDA-MB-231 or GFP MCF-7 cells were loaded in a fibrin gel in the third or bottom chamber to simulate cancer metastasis (Fig. 6A). BMoaC were cultured for an additional 8-9 days, over which time cancer cell migration was easily observed into both chambers (Fig. 6B, C).

**Figure 6.**
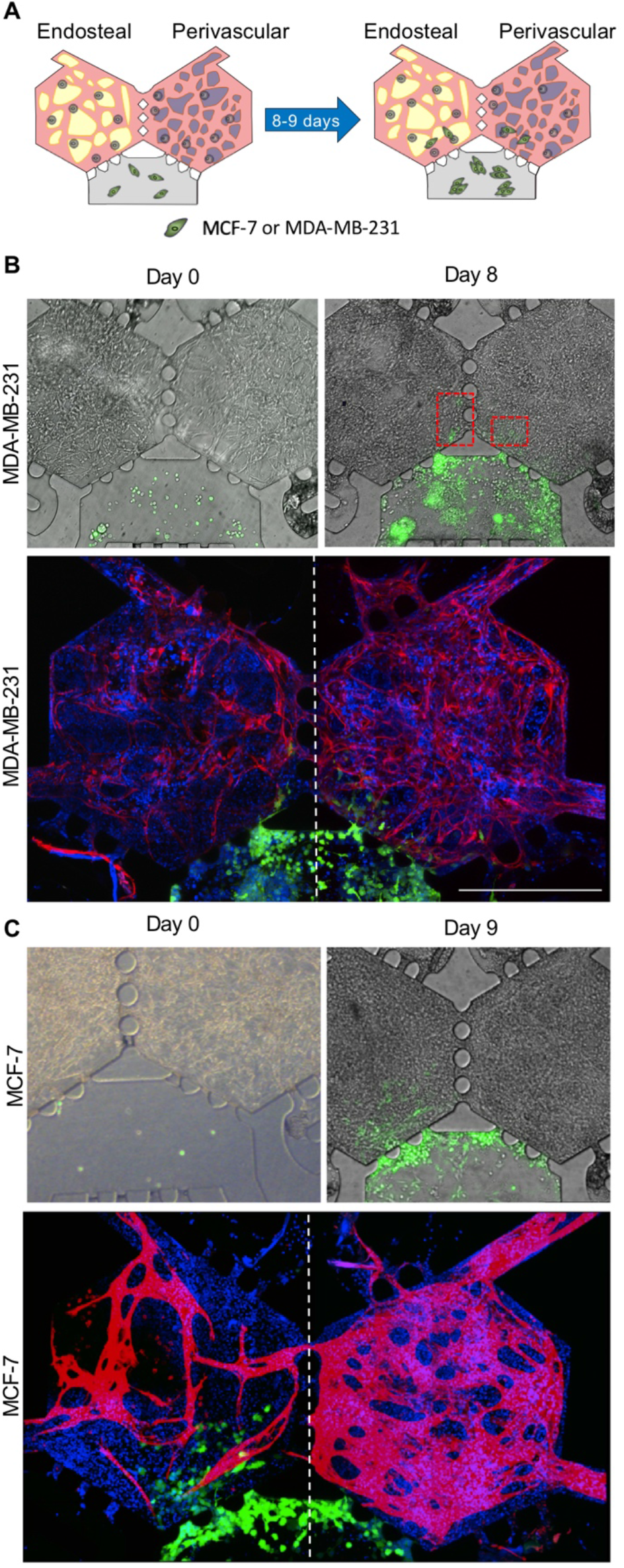
Breast cancer cells migrate toward endosteal and perivascular niches. (**A**) Experimental design: Breast cancer cells are loaded into the bottom chamber on day 4 of culture (Day 0 loading) and cultured for an additional 8-9 days. (**B**) The triple negative breast cancer cell line MDA-MB-231 (green) loaded into the device on day 0, proliferate over time and migrate into the bottom portion of both the endosteal and perivascular niche by Day 8. Red: CD34^+^ Endothelial Cells. Blue: Nuclei counterstained with DAPI. Scale bar: 500 μm. (**C**) The ER^+^ breast cancer MCF-7 cell line migrates towards the endosteal and perivascular niches after 8 days of culture in the BMoaC. Red: CD34^+^ Endothelial Cells. Blue: Nuclei counterstained with DAPI. Scale bar: 500 μm.

## Discussion

The ability to model human bone marrow function *in vitro*, such as hematopoiesis, at high spatial and temporal resolution has the potential to impact how we study and understand an array of normal biological processes in the bone marrow as well as disease. The primary goal of our platform was to create a microphysiological mimic of the bone marrow and demonstrate maintenance of the HSPC in culture, and differentiation of the HSPC into the primary lineages (i.e., hematopoiesis). Because distinct niches exist in the bone marrow (or are thought to exist), we sought to separate the perivascular and endosteal niches physically in our model system to determine if they have differential abilities to support HSPC maintenance and function. To accomplish this goal we created a model system with a dynamic 3D perfusable vascular network and separate, but adjacent, perivascular and endosteal niches. Our data demonstrates the capability of the model to maintain HSPC *in vitro* for up to two weeks independent of the niche, differentiation of the HSPC into the myeloid and erythroid lineages, intravasation of immature neutrophils into the adjacent microfluidic lines, microvascular networks with *relevant* permeability, and expression of soluble and fixed proteins consistent with the bone marrow microenvironment. Finally, our device incorporates an additional chamber that can be used to study the trafficking of tumor cells into the adjacent perivascular and endosteal niches which could provide insight into cancers that preferentially metastasize to bone marrow including breast and prostate.

Maintaining, and in particular expanding, the HSPC *in vitro* has proven challenging due to rapid differentiation and population outgrowth of cells that secrete soluble factors which inhibit HSPC expansion. Some success has been achieved with suspension and fed-batch culture systems which specifically inhibit these signals and/or stimulate the HSPC with factors such as stem regennin 1 (SR1), Notch-Delta like ligand, Angiopoietin-like 5, and IGFBP2 (*33–35*). The primary goal of these studies was expansion of the HSPC for potential clinical applications such as bone marrow transplantation. In contrast, the goal of our platform was to recreate *in vitro* an *in vivo* niche capable of maintaining a small population of HSPC. This niche is thought to be closely associated with the abluminal surface of the endothelial cell (*3, 4, 36*), and also thought to be a preferred location for metastatic breast cancer cells (*1, 13, 16*). Both of the *in vitro* niches in our platform contain an endothelial abluminal surface and both niches supported the HSPC. When devices were cultured in the absence of endothelial cells, the fibrin hydrogel was degraded by the remaining cell types (data not shown).

Previous attempts to recreate human bone marrow using tissue engineering or “organ-on-a-chip” technologies have been few, but have demonstrated success in recreating several seminal features of bone marrow including hematopoiesis, maintenance of the HSPC, the potential for breast cancer cell metastasis, and drug response (*17, 18, 37, 38*). The studies by Torisawa and colleagues (*38*) were limited by the use of mouse HSPC and *in vivo* culture time within the mouse. Sieber and colleagues (*37*) demonstrated success using human cells, but did not include endothelial cells or any model of the vascular system. Marturano-Kruik and colleagues (*38*) presented a model of bone marrow that included a bone scaffold with EC and mesenchymal stem cells. This model demonstrated the formation of rudimentary blood vessels and the potential for studying the metastasis of breast cancer cells, but did not demonstrate hematopoiesis. Most recently, Chou and colleagues (*18*) demonstrated a model that could recapitulate hematopoiesis, bone marrow drug toxicity, and features of a genetic disorder that impacts the bone marrow. However, this model was not able to demonstrate the functional maintenance of the HSPC (lack of CFU-GEMM). In addition, all of the previous models are relatively large in size (order 3-7 mm) limiting the possibility of higher throughput applications. While it is difficult to recreate all features of bone marrow in a single model, our platform demonstrates a 3D perfusable vascular network, separate perivascular and endosteal niches, hematopoiesis, migration and egress of neutrophils, the maintenance of the HSPC including the ability to form CFU-GEMM, and migration of breast cancer cells into the niche all within a size that is < 1 mm.

The two niches in our model demonstrated several similarities. For example, after 14 days of culture, both niches were able to maintain a small population of CD34^+^/CD133^+^ cells similar to adult marrow (0.1%-5%) (*39, 40*). In addition, by flow cytometry and immunofluorescence both niches were able to stimulate the differentiation of CD34+ HSPC into Lin^+^ cells, and these cells were able to exit the device from both niches and enter into the fluidic lines. This latter step mimics the *in vivo* process of leukocyte egress from the bone marrow into circulation. Thus, this platform may provide a tool for researchers to study the innate immune system *ex vivo*. Finally, tissues formed with both the osteoblast and BMSC expressed hallmark proteins and markers such as SCF, SDF-1/CXCL-12, and E-selectin/CD62E that are constitutively expressed in native bone marrow (*4, 16*). The niches displayed differences primarily in the vascular network. For example, the vasculogenic potential of the two stromal populations differed in their ability to regularly form robust vascular networks. The osteoblast cell line did not form patent vascular networks with the same reliability as the BMSC. For example, the networks derived in the presence of the osteoblast had a significantly higher number of blood vessel endpoints, indicative of a less connected network. Because networks formed with the osteoblast did not reliably form anastomosis with the fluidic lines, permeability measurements were limited to a few patent networks

As culture time increased, the number of cells capable of generating lineage-restricted (erythroid or myeloid) colony forming units appeared to increase (i.e., comparing Day 10 or 14 to Day 7) when cells were isolated from devices treated with G-CSF and IL-3. However, this was significantly different from the control only at Day 10, and the assay displayed significant variance. One limitation of the model is the small volume of the tissue chambers which limits the number of HSPC that are initially seeded. The HSPC rapidly differentiate, and thus the total number of cells that appear in each chamber is strongly dependent on the initial seeding density. The result is a wide range of dilution by differentiated cells at later timepoints. The methocult assay seeds a constant number of cells from each device/experiment, and, although we accounted for the dilution by differentiated cells, we observed a wide range of dilution factors. Thus, the small number of HSPC per chamber could account for the large variation in the number of colony forming units that we observed in the methocult assay (Fig. 4 and 5).

Besides providing a model of hematopoiesis, our microfluidic model of human bone marrow provides an additional chamber to investigate cell migration and/or soluble mediator communication between the bone marrow and an alternate tissue type. Here, we used this chamber to introduce breast cancer cells after the bone marrow had developed. We were able to observe breast cancer cells migrate into the bone marrow as well as small cells derived from the bone marrow enter into the bottom chamber and interact with the cancer cells. The platform could also be adapted for investigating hematological-based malignancies such as leukemia and myelodysplastic syndromes.

In this study, we present a novel microfluidic device that is capable of supporting two microenvironments or niches of human bone marrow, while maintaining physiological proximity as well as communication. The microtissue model of human bone marrow displayed a perfusable 3D vascular network, as well as the expression of ECM proteins and other biomarkers that are present in the native bone marrow. Importantly, CD34^+^ HSPC were cultured and maintained in the device for up to two weeks and were able to differentiate into immature neutrophils that could egress from the marrow and into adjacent microfluidic lines. Finally, an adjacent chamber can be loaded with an alternate cell of interest at a later time point to investigate interaction and communication with the bone marrow. The platform offers superior optical clarity for resolving location-dependent phenomena, such as resolving the stem cell niche and visualizing cancer metastasis and could be used as a tool for drug discovery; or be adapted to investigate hematological-based diseases.

## Materials and Methods

### Device Design and Modeling

The microfluidic device consists of two hexagonal shaped chambers connected by three symmetric two-way ports (Fig. 1A). The geometry of the port is designed using the concept of a capillary burst valve, and thus enables sequential loading of fibrin hydrogels without leaking into the adjacent chamber (*20, 21*). The port measures 80 μm in length, while each hexagonal chamber is 865 μm across, providing ample area to simulate a small number of adjacent niches in the native bone marrow. The diffusion of soluble signaling molecules as well as migration of cells between the two chambers is also enabled by the port design. A third chamber, adjacent to the hexagonal chambers, enables loading of a third cell type (e.g., cancer cell) at a later time point (Fig. 1A, B). The third chamber connects to the first two by an asymmetric one-way valve designed to prevent the fibrin gel from entering the chamber during the initial loading.

To model the molecular transport in the device, the finalized device design was imported into COMSOL Multiphysics 5.2 to simulate steady state interstitial fluid flow and concentration profiles. The fluid flow was modeled using the *transport through porous media* with a no-slip boundary condition at the walls. The tissue chambers were filled with acellular fibrin gel. The hydrostatic pressure difference created by maintaining differential levels of fluid in the inlets and outlets drove fluid through the tissue (Fig. S1), We chose to analyze transport of 70 kDa dextran molecules in the device, as most growth factors and chemokines have MW within the same order of magnitude. This transport was modeled by coupling the *transport of diluted species* and the *transport through porous media* modules. The influx and outflux conditions were matched with the actual experimental conditions. The following physical constants were used: diffusion coefficient of dextran 7 ⊠ 10^−11^ m^2^/s, porosity and hydraulic permeability of fibrin gel, 0.99 and 1.5 ⊠ 10^−13^ m^2^, respectively (*20, 22*)

### Device Preparation

A master mold of SU-8 on a silicon wafer was prepared using standard soft photolithography techniques, as previously described (*23, 24*). Briefly, a 100 μm layer of SU-8 3050 was spun onto a Si-wafer. Then, a single mask photolithography step patterned the microfluidic design onto the wafer. After silanizing the mold, polydimethylsiloxane (PDMS) was mixed with curing agent (Dow Corning) in a 10:1 ratio and poured onto the mold after degassing. The molds with PDMS were then left overnight in an oven for the PDMS to cure. The cured PDMS was removed from the mold and plasma bonded to glass cover slip (Fisher Scientific) or a glass slide (Fisher Scientific) to complete the device. Devices were exposed to ultraviolet (UV) light for at least 30 minutes to ensure sterility before loading.

### Dextran Diffusion and Fluorescence Recovery after Photobleaching (FRAP)

A fibrin gel (10 mg/mL) was injected into all three chambers of the device and allowed to polymerize for 30 minutes at 37°C. Fluidic lines were coated with 1% gelatin which was replaced by EGM-2 (Lonza) for up to 24 hours. Before imaging, EGM-2 was removed and either 70 kDa TRITC or FITC Dextran was introduced into the top fluidic lines. Diffusion through the fibrin gel was captured every five minutes for 60 minutes, totaling a series of 13 images. For FRAP experiments (performed on an Olympus FV3000), 70 kDa FITC dextran was loaded into the top port under pressure gradients of 11.5 and 20 mm H_2_O. A circular region was photobleached for 2 seconds at various points in the device and 20 images were acquired at a rate of 0.56 or 0.76 s/frame. The images were thresholded, and the center of the bleached regions was found using the circle tool in FIJI. The coordinates of the centroids were noted over time and the displacement of the centroid over time was calculated to find the interstitial velocity through the fibrin gel.

### Cell Culture

Endothelial colony-forming cells (EC) were isolated from cord blood as previously described by our group and others (*23, 25, 26*). All cord blood samples were collected in accordance with the University of California, Davis and Washington University in St. Louis School of Medicine Institutional Review Board regulations. Briefly, EC were cultured on 1% gelatin in EGM-2. All cells were used between passage 5-7, and were maintained at 37.5°C in a 5% CO_2_ incubator.

HSPC (CD34^+^) were isolated from cord blood using the EasySep™ Human Cord Blood CD34 Positive Selection Kit II (StemCell Technologies) according to the manufacture’s guidelines. Cells were isolated and frozen for future use. Cells were thawed the day of use and kept in StemSpan SFEM II media (StemCell Technologies) with CD34 supplement (StemCell Technologies) and 1 μM of StemRegenin (StemCell Technologies) before loading into the microfluidic device. The efficiency of the magnetic bead sort was assessed by flow cytometry on a small aliquot of cells. Some samples were loaded with 1 μM of CellTracker Green CMFDA (Invitrogen) per the manufacture’s instructions in order to quantify the number of CD34^+^ cells loaded into each chamber.

The human fetal osteoblast (hOB) 1.19 cell line (ATCC) was expanded at 33.5°C in DMEM/F12 media (Gibco) containing 10% FBS (Invitrogen) and 0.3 mg/mL of G418 (Invitrogen) before freezing. After thawing, cells were passaged a maximum of one time before use. Osteoblasts were cultured up to a total of 7 days in 37.5°C at 5% CO_2_.

Bone marrow stromal cells (BMSC, Lonza) were cultured in Myelocult 5100 (StemCell Technologies) supplemented with 1 μM hydrocortisone (StemCell Technologies), 2 mM L-glutamine (ThermoFisher), and 50 U/mL Penicillin-Streptomycin (ThermoFisher). All cells were used between passage 3-5.

MDA-MB-231 breast cancer cell line (triple negative) expressing red fluorescent protein (RFP) (Cell Biolabs) were cultured in DMEM (Gibco) with 10% FBS (Gibco), 2mM L-Glutamine (Gibco) and 1% Penicillin-Streptomycin (Gibco). MCF-7 (Cell Biolabs) breast cancer cells (estrogen and progesterone receptor positive) expressing green fluorescent protein (GFP) was cultured in the same media.

### Bone marrow-on-a-chip (BMoaC) model

Fibrinogen (Sigma) prepared in Dulbeco’s Phosphate-Buffed Saline DPBS (Invitrogen) was used at a final concentration of 10 mg/mL. To generate the perivascular niche, EC (1 × 10^7^ cells/mL) and BMSC (1 × 10^7^ cells/mL) were mixed together at ratio of 1:1. The cell suspension was then mixed with thrombin for a final concentration of 3 U/mL before loading into the device. Gels were allowed to sit for 3-5 minutes before loading the other side of the device. Similarly, to generate the endosteal niche, EC (1 × 10^7^ cells/mL) were mixed with OB (1 × 10^7^ cells/mL) at a 1:1 ratio. These cells were resuspended in fibrinogen (10 mg/mL final) and mixed with thrombin (3 U/mL final concentration) before loading into the device. Devices were carefully moved to the incubator and polymerized for 20-30 minutes, after which time the fluidic lines were coated with 1% gelatin. To investigate the hematopoietic niche, 1 × 10^6^ CD34^+^ cells/mL were mixed at a ratio of 1:10:10 with the endothelial cells and respective stromal cell (BMSC or hOB). To study cancer migration, tissues were allowed to form in the top chambers for 4 days before the subsequent loading of cancerous breast epithelial into the bottom chamber in a 10 mg/mL fibrin gel at a final concentration of 1 ×10^6^ cells/mL.

Devices were kept in culture over a period of 14 days. To encourage uniform vascular development within chambers and anastomosis with the microfluidic ports, the pressure heads were flipped every day for the first 4 days, after which, the pressures were held constant (*21, 27*). Devices received a 1:1 blend of fully supplemented EGM-2 and SFEM StemSpan II + 1x CD34 expansion + 1 μM SR1, unless otherwise noted. The permeability of the vessels was characterized with 70 kDa dextran as previously described (*20, 27*). Briefly, 5 μg/mL of 70 kDa FITC dextran was introduced to the top fluidic lines and allowed to perfuse through the device for 15 minutes before acquiring a series of time lapse images. The permeability (P) was calculated by quantifying the average background fluorescence intensity (I_b_), initial average fluorescence intensity (I_i_), and the final average fluorescence intensity (I_f_) after a time differential (Δt).The fluorescence intensities are determined across a blood vessel of a particular diameter (D), which may be expressed as:

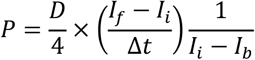

On days 7, 10, and 14, devices were assigned for either tissue analysis via immunofluorescence staining or individual cell analysis via flow cytometry and methocult colony forming unit (CFU) assays. Devices designated for staining were fixed in 10% formalin for no more than 2 hours while those designated for analysis by flow were digested with nattokinase (20 U/mL, Japan Bio Science Laboratory (*28*)). For migration studies, MDA-MB-231 or MCF7 breast cancer cells were loaded into 4-day old BMoaC in a fibrin gel at a concentration of 1×10^6^ cells/mL and kept in culture for an additional 8 days (12 days of total culture).

### Phosphorescence lifetime imaging microscopy (PhLIM)

Phosphorescence lifetime image microscopy (PhLIM) was used to measure the O_2_ tension within the device, as detailed in previous publications (*29, 30*). Briefly, our FV1200 confocal microscope (Olympus), upgraded with a phosphorescent lifetime instrument (ISS, Urbana-Champaign, IL), was modulated at 10 kHz and 5% duty cycle to excite Oxyphor G4 (O_2_ Enterprises, Philadelphia, PA) dye via a 635 nm laser. Beam emission was collected by a miniTDU with two Hamamatsu 7422p-50 detectors attached directly to the confocal head. Our confocal microscope was also installed with a stage-top incubator (Okolab, San Bruno, CA) for O_2_ and temperature control.

Experimental devices containing endosteal and perivascular niches were loaded with 20 μM Oxyphor G4 dye on days 6, 9 and 13, and O_2_ concentrations were measured on days 7, 10 and 14 at 37°C room air. A calibration curve correlating O_2_ concentration with Oxyphor G4 lifetime was generated for each device by correlating room air oxygen to lifetime measurements taken upstream of feeding lines O_2_ concentration is expected to be close to the environmental O_2_. Five repeat measurements were averaged to generate the phosphorescent lifetime of each pixel, which was then rank ordered to eliminate outliers (e.g., upper outlier is > 1.5*(IQ)+Q3 where IQ is the interquartile range and Q3 is the value of 3^rd^ quartile). The remaining data was then converted to %O_2_ using the calibration curve and a mean O_2_ percentage was generated for each tissue chamber.

### Methocult Assay

A methocult assay (MethoCult™ H4034 Optimum, StemCell^M^ Technologies) was performed per the manufacturer’s instruction, with some modification. Briefly, cells isolated from the device after nattokinase digestion were resuspended in IMDM (Gibco) with 2% FBS (Gibco) at a concentration of 1.3 × 10^3^ cells/mL. This cell suspension was mixed with MethoCult media and plated in 60 mm^2^ Gridded Scoring Dishes (Corning) such that there were 360 cells per plate. Cells were cultured for 10 days to enable colony formation before imaging on an IX83 inverted microscope (Olympus). Colony types were identified manually by blinded researchers by comparing to gold-standard images provided by the vendor. To account for varying seeding density in every trial and the respective varying proliferation rates, we normalized our cell counts for each trial and condition by comparing day 10 and day 14 to the respective day 7 CD45 population. We refer to this correction as the dilution factor (DF).

### Immunofluorescence

Devices were blocked overnight with 0.5 M Glycine (Sigma Aldrich) and 2% bovine serum albumin (BSA, Sigma Aldrich) in DPBS. Devices were stained overnight in 0.1% BSA, with all antibodies being delivered through the fluidic lines. All antibodies were chosen due to their presence in the native bone marrow and (Table S1) were used at 1:100 unless otherwise noted. Devices stained for osteopontin and leptin were permeabilized with 0.1% Tween-20. Devices were rinsed with DPBS between primary and secondary antibodies. Nuclei were counterstained with DAPI at a final concentration of 5 μg/mL or SytoxOrange (Invitrogen) at 5 μM. Devices were rinsed before imaging and incubated with ProLong™ Antifade (ThermoFisher) per the manufacture’s instructions to reduce photobleaching. Fluorescent images and timelapse were acquired with an IX83 (Olympus), while multi-area confocal images of the devices were collected with an FV3000 Confocal (Olympus). Image post-processing was done either with FV310 or in FIJI. Device images were stitched using the FIJI pairwise stitching plug-in (*31*) or by hand in PowerPoint. CD34^+^ cell quantification was done in FIJI by first cropping the image to the hexagonal chamber, thresholding the image, utilizing the watershed tool, and then analyzing particles that were greater than 100^2^ pixels and a circularity of 0.1-1.

### Flow cytometry

After 7, 10, or 14 days, BMoaC devices were digested using 20 unit/mL nattokinase (Japan Bio Science Laboratory). Devices were pooled and then placed in a gel coated well for 1 hour to deplete stromal cells from the sample. Individual cells were prepared for analysis by flow cytometry. Cells were blocked in 100 μL of a 0.1% BSA solution for 15 min at 4°C and then stained with a panel of conjugated antibodies for specific target antigens (provided in Table S2) for 1 hour at 4°C. Cells were washed with PBS and then exposed to a fixable live/dead marker for 30 min. Cells were then washed again in PBS and analyzed using an Attune NxT Acoustic Focusing Cytometer (ThermoFisher #A24858). Compensation controls were generated using single-stained populations of positive beads and negative beads (Spherotech #CMIgP-30). Data analysis and compensation correction was performed using FlowJo software. Gating of target cell populations was determined based on fluorescent minus one (FMO) controls for each fluorescent color using cryopreserved peripheral blood mononuclear cells (PBMCs).

### Statistical Analysis

All statistical analysis was performed using GraphPad Prism 8. T-tests were performed where indicated. Analysis of variance (ANOVA) followed by Tukey’s test for post-hoc comparisons were otherwise performed. All data is represented as the mean with the standard deviation included.

## Acknowledgments

### General

The authors would like to thank the Lewis Lab for use of the Invitrogen Attune NxT Flow Cytometer.

### Funding

This work was supported by the following grants from the National Institutes of Health (F32CA210540-01A1, D.E.G.; UH3TR000481, S.C.G; R21CA208519, S.C.G., K.W., and D.L.).

### Author Contributions

D.E.G. Designed the experiments, performed the experiments, data analysis and statistal analysis, and wrote the manuscript. M.B.C., P.A.S., Z.A.R., B.S.S., A.A., A.M.D., L.A., and J.M.L. performed experiments, data analysis and statistal analysis. N.R.N. performed the experiments and data analysis. K.W. and D.C.L. helped conceptualize, review and edit the manuscript. S. C. G. Designed the experiments and wrote the manuscript.

### Data and Materials Availability

All data is presented in the manuscript. Authors may be contacted for additional data related to this paper.

## Supplementary Materials

**Figure S1.**
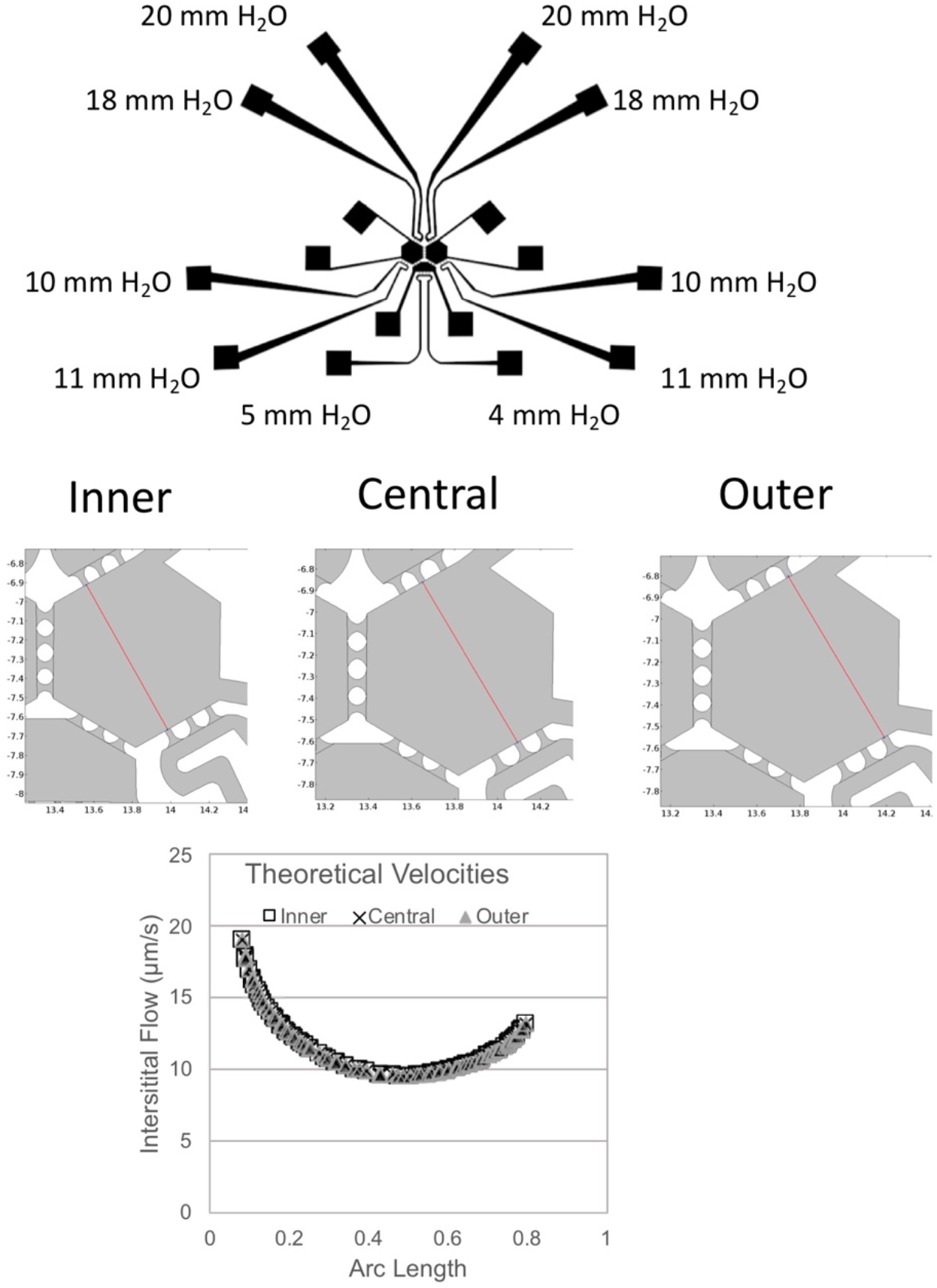
Schematic of the device depicts the pressure heads that were used for each microfluidic port. Screen-shots of three arbitrary lines draw through the ports in COMSOL were used to calculate the interstitial flow through a simulated fibrin gel in the device. When graphed, interstitial velocities through the ports show little difference.

**Figure S2.**
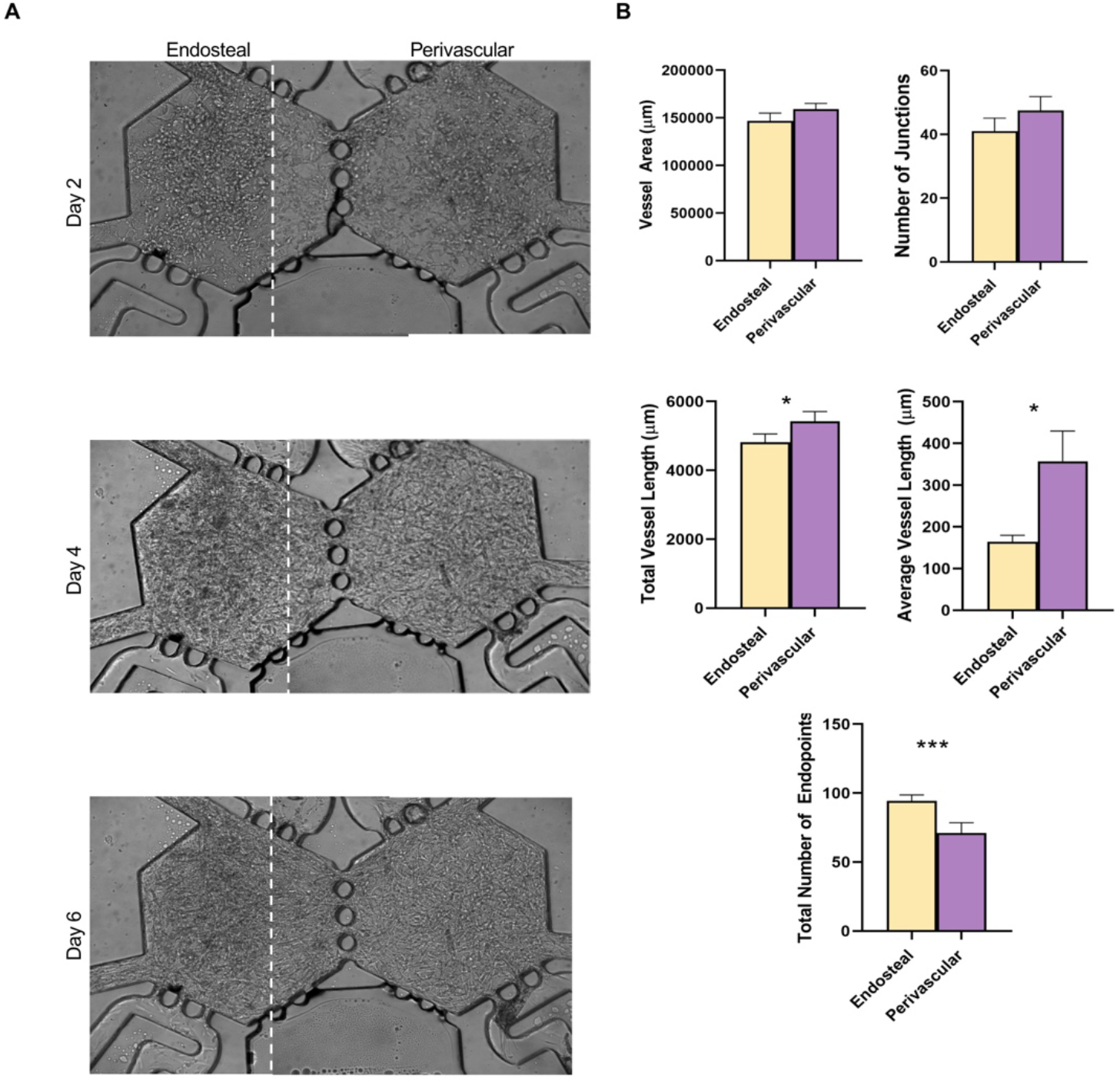
(**A**) Microvascular networks self-assemble from a cell suspension (Day 2) into vascular fragments (Day 4) and an interconnected network (Day 6). (**B**) Vascular networks stained for either CD34 or CD31 in Fig. 2 and Fig. 3 were characterized using Angiotool, Mean with SEM shown for each index; n=10. *p<0.05, **p<0.01, paired t-test.

**Figure S3.**
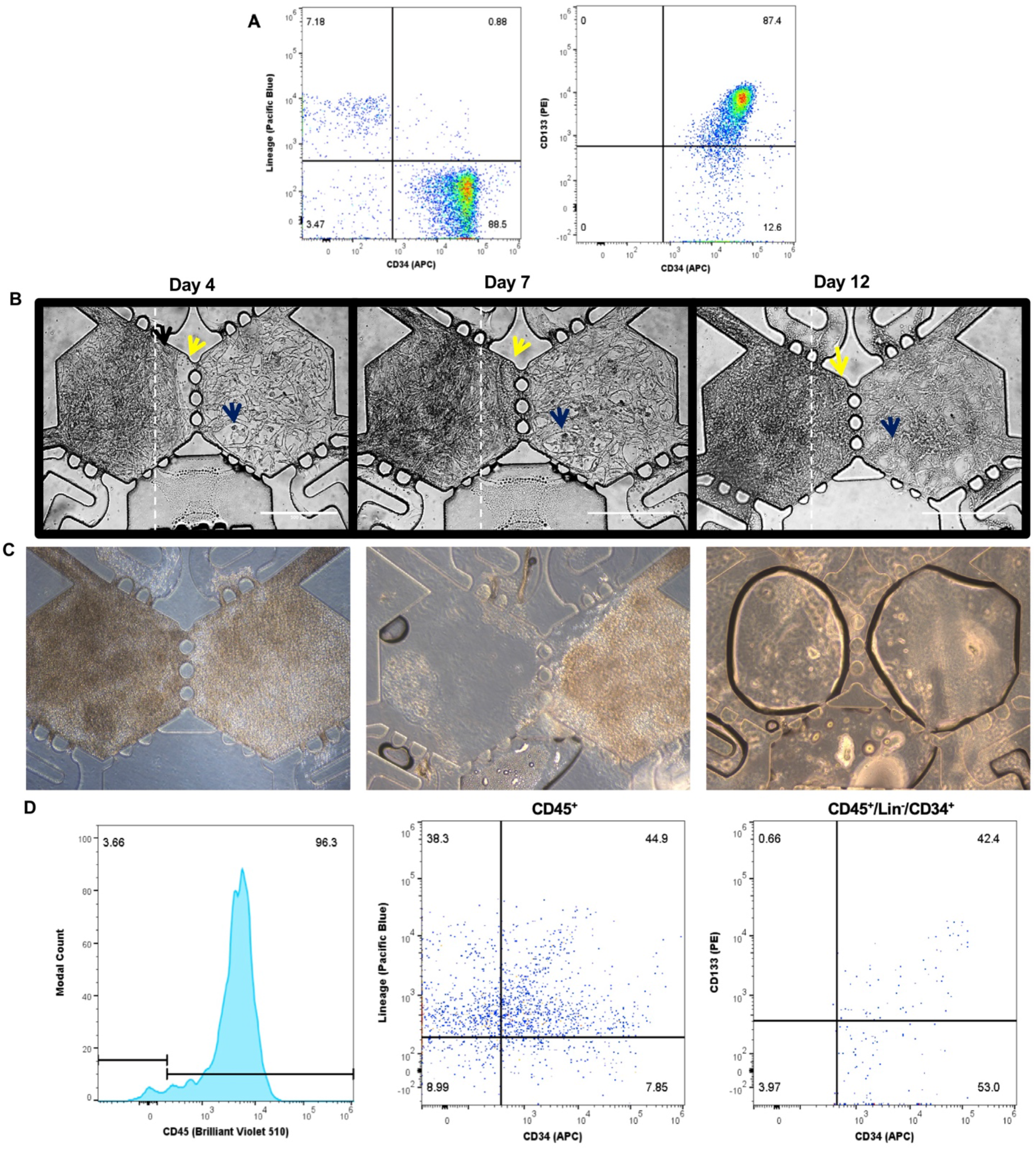
**(A)** Representative flow cytometry plot of Lin^−^/CD34^+^ and CD133^+^/CD34^+^ cells isolated from cord blood before being placed in the device.(**B**) CD34^+^ Cells proliferate in the device overtime. Yellow and dark blue arrows highlight CD34-derived cells that are proliferating in the endosteal and perivascular niches, respectively. (**C**) Representative image of a device digested with 10 U/mL nattokinase demonstrates the ability to separate the niches for anlaysis. (**D**) Representative FACS plot of the samples digested on day 10 collected from the perivascular niche.

**Figure S4.**
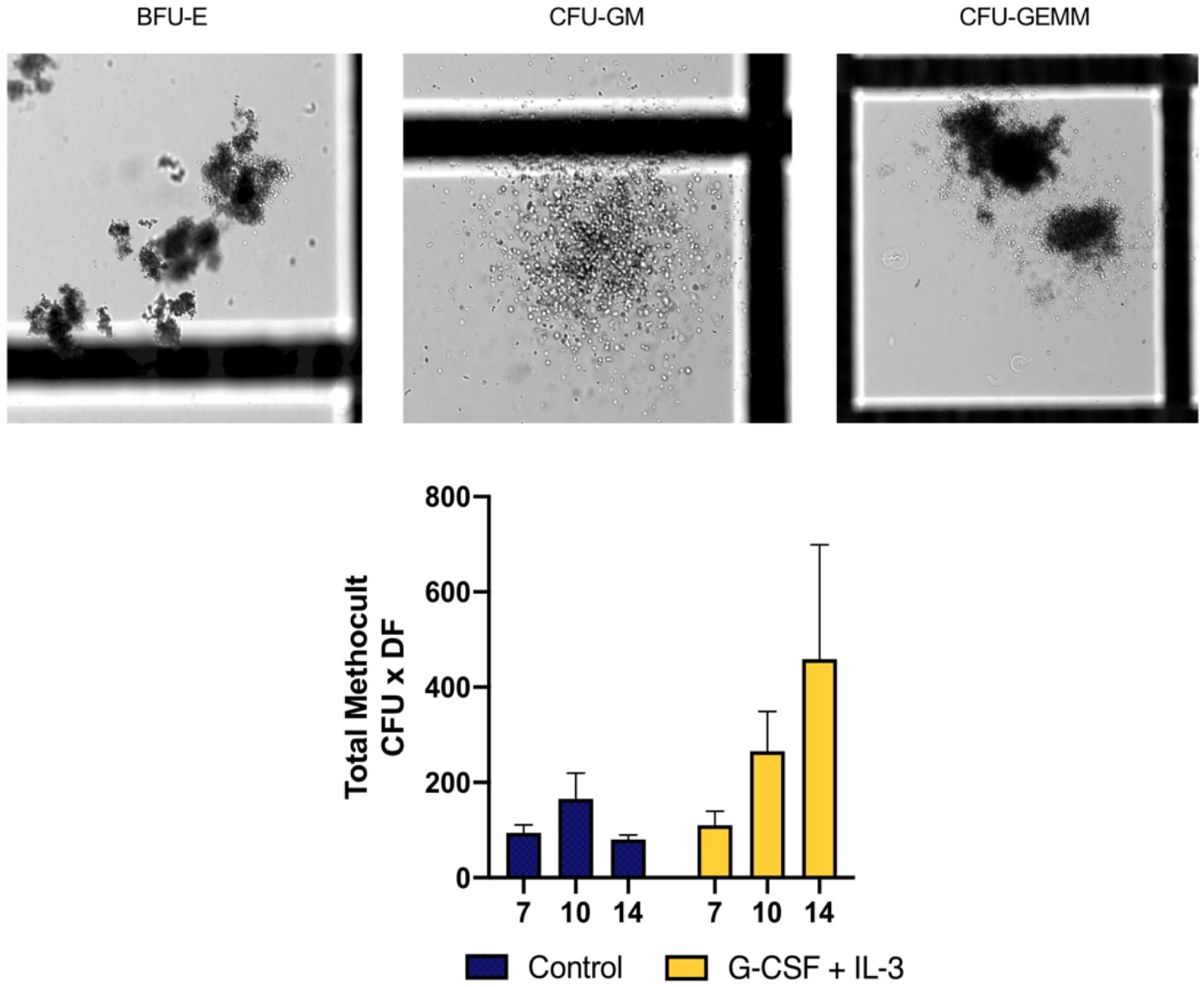
Representative images of CFU colonies generated from devices cultured for 14 days before digests. Digested cells were then cultured for an additional 10 days (24 days of culture total) in methocult before quantification. The total number of colonies generated in control conditions or when devices were stimulated with 30 ng/mL of G-CSF and IL-3, each. Mean with SEM shown, n=6.

**Figure S5.**
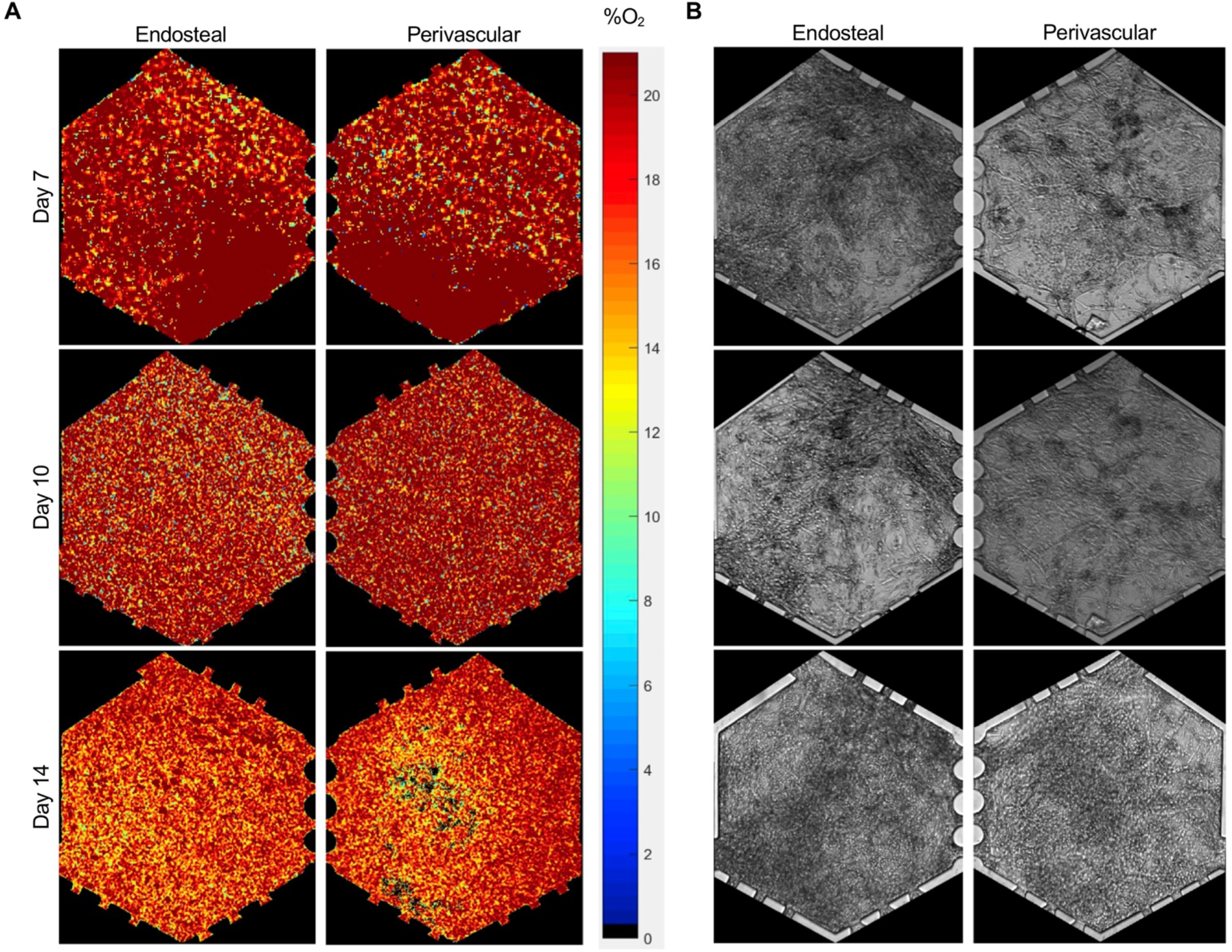
O_2_ tension within endosteal and perivascular chambers decreases over time. (**A**) O_2_ tension maps of representative endosteal and perivascular chambers from the same device is shown for days 7, 10, and 14. (**B**) Bright field images of tissue chambers from A.

**Table S1:** Immunofluorescent Antibodies

| *Antigen* | *Fluorophore/host* | *Vendor* | *Cat #* |
| --- | --- | --- | --- |
| CD34 | APC | BioLegend | 343510 |
| Laminin | Alexa Fluor 488 | Novus Biologicals | NB300-144AF488 |
| CD54 | Alexa Fluor 488 | BioLegend | 322713 |
| CD62E | APC | BD Biosciences | 551144 |
| CXCL-12 | PE | R&D Systems | IC350P |
| Nestin | Alexa Fluor 488 | ThermoFisher | 53-9843-80 |
| CD90 | FITC | BioLegend | 328108 |
| CD14 | Brilliant Violet 421 | BioLegend | 325628 |
| Lineage | Pacific Blue | BioLegend | 348805 |
| CD31 (JC/70A) | Mouse | ThermoFisher | MS-353-S |
| VCAM-1 | Rabbit | Abcam | ab134047 |
| Osteopontin* | Rabbit | Abcam | ab8448 |
| Leptin | Rabbit | ThermoFisher | PA1-052 |
| Goat anti-Rabbit IgG* | Alexa Fluor 488 | Invitrogen | A-11008 |
| Goat anti-Mouse IgG* | Alexa Fluor 555 | Invitrogen | A-21422 |
| SytoxOrange | N/A | ThermoFisher | S11368 |
| DAPI | N/A | ThermoFisher | D1306 |

\*Used at 1:1000

**Table S2:** FACS Antibodies 2

| *Antigen* | *Fluorophore* | *Vendor* | *Cat #* | *Vol. per test* |
| --- | --- | --- | --- | --- |
| CD14 | Alexa Fluor 700 | BioLegend | 325614 | 5 µL |
| CD15 | APC/Cy7 | BioLegend | 323048 | 5 µL |
| CD33 | Brilliant Violet 510 | BioLegend | 366610 | 5 µL |
| CD34 | APC | BioLegend | 343510 | 5 µL |
| CD45 | FITC | BioLegend | 368508 | 5 µL |
| CD133 | PE | BioLegend | 372804 | 5 µL |
| Lineage | Pacific Blue | BioLegend | 348805 | 20 µL |
| Live/Dead | Fixable Yellow | ThermoFisher | L34959 | 1 µL |

**Table S3:** Mean values ± SD for CFU from devices digested on day 7, 10, and 14.

|  |  | **Control**<br>**BFU-E/CFU-E** | **Control**<br>**CFU-G/GM/M** | **Control**<br>**CFU-GEMM** |
| --- | --- | --- | --- | --- |
| Day | **7** | 39.5 ± 19.1 | 44.3 ± 15.6 | 10.5 ± 7.8 |
| | **10** | 72.7 ± 52.2 | 76.4 ± 57.5 | 7.12 ± 4.7 |
| | **14** | 22.7 ± 15.2 | 49.7 ± 13.7 | 8.2 ± 4.9 |
|  |  | **G-CSF + IL-3**<br>**BFU-E/CFU-E** | **G-CSF + IL-3 CFU-**<br>**G/GM/M** | **G-CSF + IL-3**<br>**CFU-GEMM** |
| Day | **7** | 62.7 ± 40.6 | 37.8 ± 23.3 | 10 ± 8.3 |
| | **10** | 222.4 ± 60.8 | 93.7 ± 92.4 | 25.5 ± 18.4 |
| | **14** | 43.9 ± 10.7 | 215.7 ± 280.6 | 6.5 ± 8.9 |

**Movie S1.** Microvascular networks generated in the device are perfused with 70 kDa dextran.

**Movie S2**. 10-hour timelapse video with images acquired every twenty minutes shows CD34^+^-derived cells migrating through the device, into the fluidic lines, and the adjacent chamber. Images acquired on Day 6 of culture.

**Movie S3**. Confocal Z-stack of Fig. 3Ci: A blood vessel surrounds a colony of CD34^+^ and Lin^+^ cells in the endosteal niche.

**Movie S4**. Confocal Z-stack of Fig. 3Cii. CD34^+^ and Lin^+^ are seen coming in and out of focus. A blood vessel is located on the left side of the images.

